# Near-infra red light and mitochondrial large-conductance calcium-activated potassium channels: protection of hippocampal neurons, influence on channel activity and transcriptome remodelling

**DOI:** 10.64898/2026.06.09.731043

**Authors:** Piotr Bednarczyk, Małgorzata Beręsewicz-Haller, Joanna Lewandowska, Bogusz Kulawiak, Antoni Wrzosek, Barbara Zabłocka, Adam Szewczyk, Barbara Kalenik

**Affiliations:** Department of Physics and Biophysics, Institute of Biology, Warsaw University of Life Sciences SGGW, Nowoursynowska 159, 02-776 Warsaw, Poland; Molecular Biology Unit, Mossakowski Medical Research Institute, Polish Academy of Sciences, Pawinskiego 5, 02-106 Warsaw, Poland; Laboratory of Intracellular Ion Channels, Nencki Institute of Experimental Biology, Polish Academy of Sciences, Pasteura 3, 02-093 Warsaw, Poland

**Author notes:** corresponding author - Nencki Institute of Experimental Biology PAS, 3 Pasteur Street, 02-093 Warsaw, Poland. equal contribution (Piotr Bednarczyk, Małgorzata Beręsewicz-Haller).

**Keywords:** mitochondrial potassium channels, cytochrome c oxidase, near-infra red light, neuroprotection, transcriptome, hippocampus, glioma

## Abstract

Photobiomodulation (PBM) is a therapeutic approach based on illumination with red or near-infrared (NIR) light. Cytochrome c oxidase (COX), a terminal enzyme of the mitochondrial respiratory chain, contains copper centers (CuA and CuB) that absorb light within the red and NIR spectral range, making it a potential primary photoacceptor at wavelengths around 820 nm. PBM appears to be a promising strategy for the treatment and prevention of neurological disorders. Elucidating its precise molecular mechanisms may help optimize therapeutic outcomes.

Using patch-clamp method, we showed that illumination with 820 nm light activates mitochondrial large-conductance calcium-activated potassium (mitoBK_Ca_) channels in rat hippocampal mitochondria. Moreover, 820 nm light caused neuroprotective effect in NMDA-treated organotypic hippocampal cultures. Consistently, activation of mitoBK_Ca_ channel by 820 nm light illumination was observed in mitochondria isolated from glioma U-87 MG cells. To further investigate the role of mitoBK_Ca_ channel, we used CRISPR/Cas9- developed U-87 MG cells lacking the α-subunit of the BK_Ca_ channel (dBK cells). Comparative transcriptomic analysis of illuminated wild-type and dBK cells revealed significant differences in gene expression profiles. In summary, our results show two types of cellular responses to the PBM. An acute effect involving activation of the mitoBK_Ca_ channel and a long-term effect associated with extensive transcriptome remodeling. Both mechanisms may contribute to the cytoprotective effect of 820 nm near-infrared light.

**Highlights:**

- 820 nm light activates hippocampal mitochondrial BK_Ca_ channels
- 820 nm light induces hippocampal neuroprotection under excitotoxic conditions
- 820 nm light causes intensive transcriptome remodeling in glioma cells
- BK_Ca_ channels modulate a subset of transcriptomic responses to 820 nm light

**Graphical abstract:** 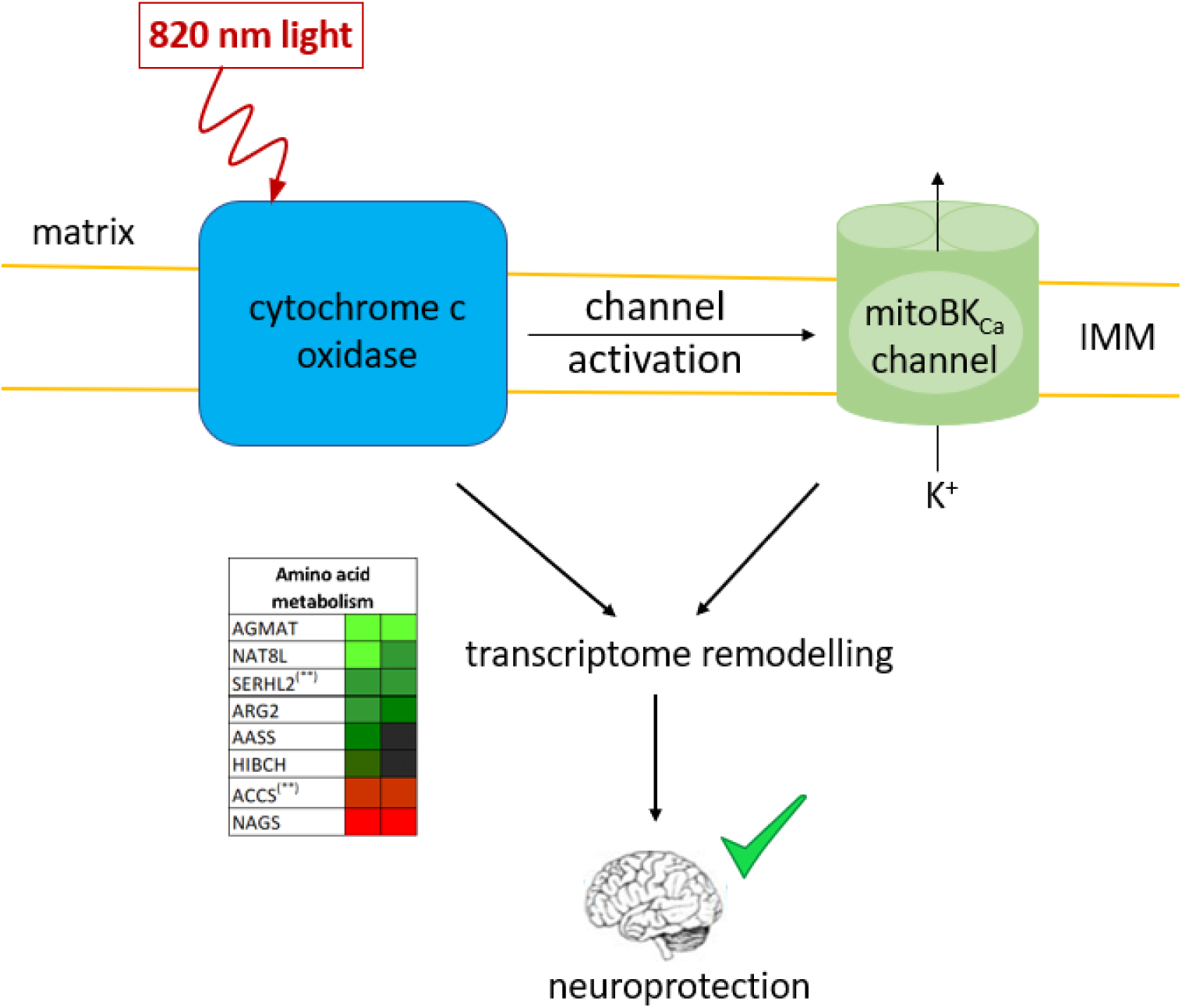

## 1. Introduction

Photobiomodulation (PBM) is a process in which beneficial effect on cells or tissues is achieved with the use of red or near-infrared (NIR) light at low power densities [1, 2]. It is a rapidly growing field with applications not only in wound healing and joint regeneration, but also in the treatment of traumatic events, degenerative diseases and psychiatric disorders [3]. There have been over 1,000 papers published on experimental laboratory studies in PBM, and another large range (1000+) of clinical studies of its effect [2]. Despite the overwhelming number of positive clinical outcomes, variations in study designs led to conflicting results including negative trials which have generated some controversy [1, 4]. Effective red and NIR light therapy is dependent on treatment parameters, thus understanding the mechanisms of action of PBM can help in parameters and results optimization [4, 5].

The main absorber of red and NIR light in the cell is cytochrome c oxidase (COX, complex IV). It is the terminal enzyme of the respiratory chain located in the inner mitochondrial membrane [2, 4, 6]. It contains two copper active centers: Cu_A_, located in subunit II, and Cu_B_, located in subunit I as part of the heme a3–Cu_B_ binuclear center.Maximum light absorption for an oxidized Cu_A_ center is observed for near-infrared light having a wavelength of 820 nm [7]. Interestingly, light around 820 nm activates COX complex, while other wavelengths can inhibit its activity [8]. PBM has been reported as beneficial through activation of COX [9]. Moreover, PBM with 820 nm light was beneficial in many conditions like treatment of chronic testicular pain [10] and orchialgia [11]; treating symptoms of Hashimoto thyroiditis [12, 13] and dental therapies [14, 15]. Furthermore, 14 days of PBM with 820 nm light was beneficial in patients with major depressive disorder and the effect remained significant at 8-week follow-up [16]. Additionally, studies using 830-850 nm and 808/905 nm improved facial nerve function recovery in Bell’s palsy patients [17] and 820 nm therapy was beneficial in myofascial pain dysfunction syndrome [18].

We have shown that COX is functionally coupled to the glioma mitochondrial BK_Ca_ (mitoBK_Ca_) channel [19]. The redox state of the respiratory chain was able to modulate mitoBK_Ca_ channel activity [19]. Recently, we observed that infrared light was able to stimulate the mitoBK_Ca_ channel in glioma cells [20]. Simultaneously, neuroprotective action of potassium channels was reported both in ischemia-reperfusion injury and in glutamate excitotoxicity [21–23]. Therefore, the aim of our study was to investigate whether the absorption of red light by COX, potentially leading to the activation mitochondrial potassium channels, can induce neuroprotective effects.

We investigated the activity of the mitoBK_Ca_ channel in mitoplasts isolated from rat hippocampi using the patch-clamp technique. Then, we examined changes in channel activity upon illumination with 820 nm infrared light. Next, we evaluated the neuroprotective effects of such illumination applied prior to excitotoxic stress in organotypic hippocampal slice cultures. Finally, we performed a comparative transcriptomic analysis of near-infrared illuminated wild-type and BK_Ca_ α-subunit knockout U-87 MG glioma cells, focusing on differential expression of genes associated with mitochondrial function. Overall, our findings point to potential molecular targets underlying the beneficial effects of infrared light in the brain.

## 2. Methods

### 2.1. Preparation of mitochondria from neonatal rat hippocampal neurons

A mitochondria-enriched fraction was isolated according to previously described protocols [24–26]. Briefly, freshly dissected hippocampi from control rats were homogenized in ice-cold mitochondria isolation buffer composed of 210 mM mannitol, 70 mM sucrose, 5 mM HEPES, 1 mM EGTA, and 0.5% (w/v) fatty acid–free BSA at pH 7.2, using a Teflon–glass homogenizer. The homogenate was centrifuged at 1,000 × g for 10 min at 4°C to remove nuclei and cellular debris. The resulting supernatant was subsequently centrifuged at 11,000 × g for 20 min at 4°C to obtain a crude mitochondrial pellet. The pellet was gently resuspended in an isolation buffer and used immediately for patch-clamp experiments.

### 2.2. Cell culture

U-87 MG wt cells were grown following a standard procedure described in [27]. U-87 MG glioma cells and derived knockout (dBK) cell line were cultured in DMEM supplemented with 10 % FCS, 2 mM L-glutamine, 100 U/ml penicillin, and 100 μg/ml streptomycin at 37 °C in a humidified atmosphere with 5 % CO_2_. The cells were fed and reseeded every third day. Cell lines lacking the α-subunit of the BK_Ca_ channel were previously generated in our laboratory [27].

### 2.3. Preparation of mitochondria from U-87 MG glioma cells

Mitochondria from U-87 MG cells were prepared as previously described [19]. Glioma cells from two culture flasks were collected in PBS medium and centrifuged at 400 × g for 10 min. The cell pellet was resuspended and homogenized in a preparation solution (250 mM sucrose, 5 mM HEPES, pH = 7.2). To isolate the mitochondria, the homogenate was centrifuged at 9,200 × g for 10 min. The pellet was then suspended and centrifuged at 780 × g for 10 min. The supernatant was transferred to a new tube and centrifuged at 9,200 × g for 10 min. Finally, the pelleted mitochondria were then resuspended in a storage solution (150 mM KCl, 10 mM HEPES, pH = 7.2) and centrifuged at 9,200 g for 10 min. In the final step, the mitochondria were resuspended in 0.1 ml of a storage solution. All procedures were performed at 4 °C.

### 2.4. Patch-clamp experiments

Patch-clamp experiments using hippocampal and U-87 MG cell mitoplasts were performed similarly to previous experiments [19, 28]. In brief, mitoplasts were isolated from mitochondria by placing them in a hypotonic solution (5 mM HEPES, 100 μM CaCl_2_, pH 7.2) for approximately 2 minutes to induce swelling and rupture of the outer membrane. A hypertonic solution (750 mM KCl, 30 mM HEPES, 100 μM CaCl_2_, pH 7.2) was added to restore isotonicity. The final bath isotonic solution contained 150 mM KCl, 10 mM HEPES, and 100 μM CaCl_2_ at pH 7.2. A patch-clamp pipette was filled with an isotonic solution. All of the modulators of the mitoBK_Ca_ channel were added as dilutions in isotonic solution. For the calcium-dependence experiments, the concentration of free Ca^2+^ was controlled with EGTA, and appropriate concentrations of CaCl_2_ were calculated with MaxChelator software58 (Stanford University, Stanford, CA, USA). To apply channel modulators and isotonic solutions with different calcium concentrations, a perfusion system was used. The mitoplasts at the tip of the measuring pipette were transferred into the openings of a multibarrel “sewer pipe” system in which their outer faces were rinsed with the test solutions. The current–time traces of the experiments were recorded in single-channel mode. The pipettes were made of borosilicate glass and had a resistance of 10–20 MΩ (Harvard Apparatus GC150-10). A PC-10 puller (Narishige) was used. The currents were low-pass filtered at 1 kHz (amplifiers: Axopatch 200B, digidata: Axon 1440A, Molecular Devices). The traces of the experiments were recorded in single-channel mode. For data analysis, Clampfit 10.7 software (Axon Instruments, Molecular Devices) was used. The conductance of the channel was calculated from the current–voltage relationship (Fig. 1B). The open probability (NPo) of the N channels was determined using the single-channel search mode. Changes in the ion current were determined by event statistics. For multichannel recordings, the area (pA*s) under the curve was calculated. The baseline was fixed individually for each experiment based on the closed state of the channels (pA). The maximum number of channels in the patch was obtained by dividing the peak amplitude (pA) by the current of a single channel (pA).

**Fig. 1.**
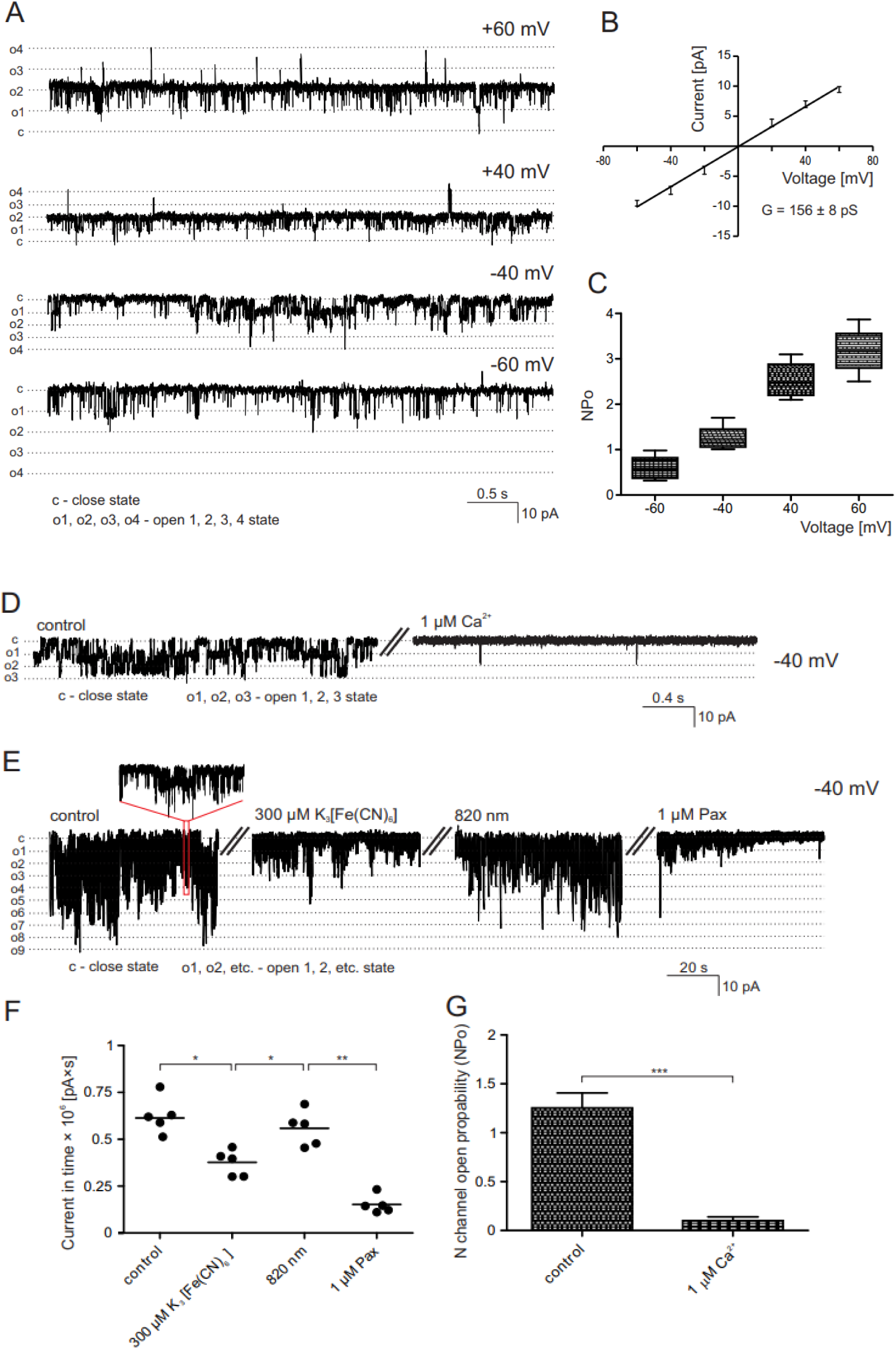
Mitoplast patch-clamp measurement and channels recordings before and after illumination. “NP_o_” represents open probability analysis of multi-channel recordings. “c” indicates a closed and “o1-o9” – open channel states. (A) Recordings of the mitoBK_Ca_ channel activity in mitoplast isolated from rat hippocampi, measured in a symmetric 150/150 mM KCl isotonic solution (in the presence of 100 μM Ca^2+^) at different voltages (60, 40, −40, −60 mV). (B) Biophysical properties of mitoBK_Ca_ channel recorded in mitoplast isolated from rat hippocampi. The current-voltage relationship, based on single-channel recordings in a symmetric 150/150 mM KCl isotonic solution (100 μM Ca^2+^) of the channel (n = 3). The conductance of the channel, as calculated based on the presented I-V curve, was equal to 156 ± 8 pS. (C) Analysis of the channel open probability (NP_o_) in control condition 100 μM Ca^2+^ at different voltages (n = 4). (D, G) NP_o_ strongly decreases in 1μM Ca^2+^. Currents were recorded at −40 mV. (E, F) NP_o_ of the mitoBK_Ca_ channel significantly decreases after admission of 300 μM K_3_[Fe(CN)_6_]. The observed effect is reversed by 820 nm light illumination and channel activity is blocked by 1 μM paxilline. Currents were recorded at −40 mV.

### 2.5. Organotypic hippocampal cultures

Hippocampal slices were prepared from 6-7 day-old Wistar rats using the slightly modified Stoppini method [29], as previously described [30–32]. Briefly, isolated hippocampi were cut into 400 µm thick slices using a McIlwain Tissue Chopper and transferred to Millicell-CM membranes (Millipore) for further growth in a humidified atmosphere of 5% CO_2_ at 36°C for 8 days. Slices were initially cultured in medium containing 50% Neurobasal (Gibco), 25% horse serum (Gibco), 22% HBSS (Gibco), 5 mg/mL glucose (Sigma), B-27 supplement (1:200, Gibco), 1 M HEPES (Gibco), and antibiotic antimitotic solution (1:100, Sigma). Then, starting from the third day of culture, 25% horse serum was gradually removed to 0%. On the eighth day of culture, using the fluorescent cell-death marker propidium iodide (PI), all slices were tested for viability to remove those showing any signs of neurodegeneration.

### 2.6. Optimization of illumination of organotypic hippocampal cultures

Hippocampal slices were illuminated with M940L3-C1 lamps (ThorLabs) modified to provide 820 nm light. Lamps were powered by Advanced Four-Channel LED Driver DC4100 with usage of DC4100-HUB (both ThorLabs). The light parameters were optimized for equal illumination of the whole culture dish ø 35 mm with power 73 mW/cm^2^. To achieve this, constant current of 50 mA was set at the LED Driver and the path of the light was established to 370 mm for the 820 nm illumination. Light intensity was measured by S120VC Photodiode Power Sensor with PM200 - Touch Screen Power and Energy Meter Console (both ThorLabs).

### 2.7. NMDA-induced excitotoxity in organotypic hippocampal cultures

Slices were treated with NIR 820 nm illumination (conditions as above described, total fluence of 525.6 J/cm^2^) or 10 µM BK_Ca_ channel activator NS1619 dissolved in DMSO for 2h, followed by excitotoxic damage induced by a 3-hour incubation with 25 µM NMDA. After 24 hours, fluorescent images were obtained using a ZEN microscope (Zeiss 510), and analyzed with ImageJ analysis software. Neuronal damage was performed by measuring PI intensity from total slice area and normalized to the maximal fluorescence intensity obtained by treating slices with 100 µM NMDA and presented as % of maximal neuronal damage.

### 2.8. RNA-Seq analysis of illuminated cells

U-87 MG wt and dBK cells were seeded to 6-well plates 4×10^4^ cells/well, 48 h prior to illumination, which was provided in a custom-made box with a stand and 6 LED830L (ThorLabs) connected in parallel. Light intensity was measured as described in the ‘illumination of hippocampal slices’ chapter and the path of the light was optimized to provide a power of 73 mW/cm^2^ in the cell culture plates. Cells were illuminated for 2h with a total fluence of 525.6 J/cm^2^.

RNA was isolated from cells with RNeasy Mini Kit (Qiagen) 30 minutes after cell illumination. RNA concentration, quality and integrity was assessed with Nanodrop One (ThermoScientific) and Agilent 2100 Bioanalyzer using an RNA 6000 Nano Kit (Agilent Technologies, Ltd.). Bioanalyzer analysis, libraries preparation, sequencing and primary data analysis was done as a service by the Laboratory of Sequencing (Nencki Institute of Experimental Biology PAS). In total, strand-specific polyA enriched RNA libraries were prepared using the KAPA Stranded mRNA Sample Preparation Kit according to the manufacturer’s protocol (Kapa Biosystems, MA, USA). Briefly, mRNA molecules were enriched from 500 ng of total RNA using poly-T oligo-attached magnetic beads (Kapa Biosystems, MA, USA). Obtained mRNA was fragmented and the first-strand cDNA was synthesized using a reverse transcriptase. Second cDNA synthesis was performed to generate double-stranded cDNA (dsDNA). Adenosines were added to the 3′ ends of dsDNA and adapters were ligated (adapters from NEB, Ipswich, MA, USA). Following the adapter ligation, uracil in a loop structure of adapter was digested by USER enzyme from NEB (Ipswich, MA, USA). Adapters containing DNA fragments were amplified by PCR using NEB starters (Ipswich MA, USA). Library evaluation was done with Agilent 2100 Bioanalyzer using the Agilent DNA High Sensitivity chip (Agilent Technologies, Ltd.) Mean library size was 300bp. Libraries were quantified using a Quantus fluorometer and QuantiFluor double stranded DNA System (Promega, Madison, Wisconsin, USA). Libraries were paired-end sequenced (2×151bp) on NovaSeq 6000 (Illumina, San Diego, CA 92122 USA).

### 2.9. RNA-Seq data preprocessing and analysis

Quality control of reads was performed using FastQC (v. 0.11.9). Adapter and quality trimming was performed using cutadapt (v. 3.4) and TrimGalore (v. 0.6.7), respectively. TrimGalore quality parameter was set to 25. Alignment of reads to the human reference genome (GRCh38) was performed using STAR (v. 2.7.9a) with default settings. The GTF annotation file used was from the 105 Ensembl release. Duplicate reads were marked using Picard MarkDuplicates (v. 2.27.4-SNAPSHOT). Final quality control was collated with MultiQC (v. 1.13) from RSeQC (v. 3.0.1) and the tools described above. Reads were summarized and counted by featureCounts (v. 2.0.0) on paired-end reads, with only primary alignments and reversely stranded reads counted. The minimum mapping quality score required for a read to be counted was set to 3. The differential and functional analyses were performed in the R environment (v. 4.1.3). The differential analysis was performed using DESeq2 (v. 1.34) with default parameters. If the sum of counts for a certain feature was less than 10 in all samples, this feature was discarded from the analysis. The functional analysis was performed using the enrichPathway() function and the gene set enrichment analysis was performed using the gsePathway() function, both from the ReactomePA package (v. 1.38). P-value < 0.05 was considered significant. Additionally grouping of enriched pathways was performed on the basis of pathways description on the reactome.org website. Volcano plots were generated using the ggplot2 package (v. 3.3.6). Venn diagrams were prepared with an online tool from: https://bioinformatics.psb.ugent.be/webtools/Venn/ website. Charts describing pathways and heatmaps were prepared with MS Office 16 suite. List of differentially expressed genes (DEGs) were screened for DEGs encoding proteins important for mitochondrial functioning with Human MitoCarta3.0 dataset available from Broad Institute website (https://personal.broadinstitute.org/scalvo/MitoCarta3.0/human.mitocarta3.0.html).

## 3. Results

To determine whether near-infrared light influences mitoBK_Ca_ channel activity and downstream cellular responses, we combined electrophysiological, neurobiological, and transcriptomic analyses. The effects of 820 nm illumination were first examined at the level of single mitoBK_Ca_ channels in isolated mitochondria, followed by assessment of neuroprotection in organotypic hippocampal cultures and transcriptomic profiling of wild-type and BKCa-deficient U-87 MG cells.

### 3.1. Electrophysiological characterization of mitoBKCa channels and their modulation

To determine whether activation of COX complex by 820 nm light modulates mitoBK_Ca_ channel activity, we first performed patch-clamp analysis of rat hippocampal mitoplasts.

The initial set of recordings was aimed at identifying the channel observed in rat hippocampal mitoplasts. These experiments showed channel activity typical for large conductance Ca^2+^-activated K^+^-channels (BK_Ca_ channels). Open probability (NPo) of the channel, recorded from −60 to +60 mV positively correlated with voltage (Fig. 1A-C) and was significantly reduced at 1µM Ca^2+^ (Fig. 1D, G). Moreover, 1µM paxilline – a selective mitoBK_Ca_ channel blocker, significantly decreased NPo, confirming channel identity.

Next, in order to standardize the redox state of the respiratory chain we used potassium ferricyanide (K_3_[Fe(CN)_6_]), which was applied during patch-clamp recordings of channel activity. Oxidation of cytochrome c oxidase (COX) by K_3_[Fe(CN)_6_] decreases activity of the mitoBK_Ca_ channels (Fig. 1E, F). Subsequent illumination of the patch with 820 nm infrared light, corresponding to the maximum absorption of the Cu_A_ chromophore of COX, restored mitoBK_Ca_ channels activity (Fig. 1E, F). These findings demonstrate that mitoBK_Ca_ channel activity is modulated by the redox state of COX and can be restored by NIR targeting this respiratory chain complex.

### 3.2. Neuroprotective effects of NIR illumination in organotypic hippocampal cultures

In the subsequent part of the study, organotypic hippocampal cultures were used to evaluate the effect of infrared light on tissue viability under stress conditions. As is shown in Fig. 2A, organotypic hippocampal cultures were subjected to 2 hours of 820 nm illumination or treated with NS1619, a BK_Ca_ channel activator, followed by 3h NMDA exposure. Neuronal viability within the hippocampus was assessed 24 hours later based on the intensity of propidium iodide fluorescence and expressed as a percentage of maximal damage. As shown in Fig. 2B, 3-hour incubation with NMDA resulted in approximately 80 % of maximal neuronal damage (81,7 ± 6,8 SD, n = 5, p<0,001 vs control). Pre-treatment with NS1619 significantly reduced NMDA-induced injury to 47.76 ± 9.63 % (∼42 % reduction, n = 10, p<0.01 vs NMDA), consistent with our earlier findings [26]. NIR illumination also significantly attenuated neuronal damage following excitotoxic insult, reducing it to 63.15 ± 11.60 % (∼23 % reduction, n= 7, p < 0.05 vs. NMDA). Importantly, neither 820 nm illumination nor NS1619 or its solvent at the applied concentration affected the viability of organotypic hippocampal cultures under control conditions (Fig. 2B). These results indicate that NIR illumination confers neuroprotection when applied prior to excitotoxic injury.

**Fig. 2.**
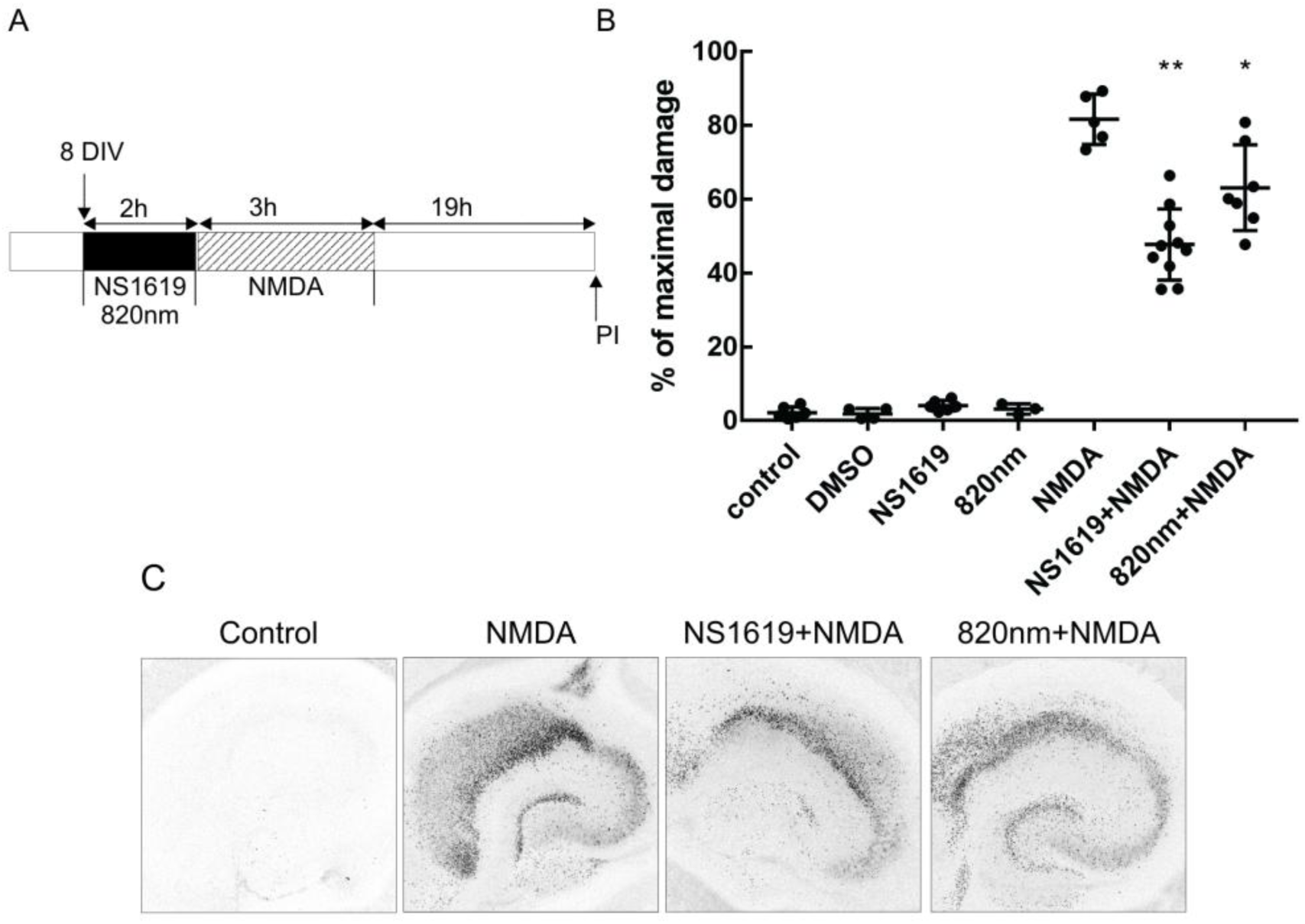
The effect of NIR on the survival of neurons treated with NMDA in organotypic hippocampal culture (OHC). (A) OHCs were cultured for 8 days to be followed by 2 hour administration of NS1619 (10 µM) or 820 nm illumination preceding 3-hour incubation with 25 µM NMDA. Neuronal damage was assessed 24 hours later using propidium iodide-staining and shown in (B) as a percent of maximal neuronal damage. Results are expressed as mean with SD, n≥3, *p<0.05, **p<0.01 versus NMDA. (C) Colour-inverted fluorescent images of propidium iodide-stained hippocampal slices 24 hours after administration of 25 μM NMDA alone or preceded by administration of NS1619 or 820 nm illumination.

### 3.3. Redox and NIR-dependent regulation of mitoBK_Ca_ channels in glioma cells

The above experiments indicate that mitoBK_Ca_ activity is regulated by cytochrome c oxidase in response to infrared light stimulation. However, the next question was which signaling pathways are activated by irradiation and the resulting activation of the mitoBK_Ca_ channel. To address this, we aimed to analyze changes in gene transcription using RNA-seq. To follow the 3R principle we replaced animal-derived material with glioma U-87 MG cell line in these experiments. We found this replacement possible if patch-clamp results obtained in this model were consistent with those described in 3.1 chapter. The basal activity of the mitoBK_Ca_ channel in U-87 MG cells (Fig. 3A) has been previously characterized by our group [19, 27, 33]. Similar to hippocampal mitoplasts, channel activity recorded at −40mV in U-87 MG cell mitoplasts was significantly reduced after admission of 300 μM K_3_[Fe(CN)_6_] and subsequently restored after 820 nm light illumination (Fig. 3B, C). These findings indicate that mitoBK_Ca_ channel activity is regulated in a manner similar to that observed in hippocampal mitoplasts. Accordingly, these results confirm the suitability of the U-87 MG cell model for further studies.

**Fig. 3.**
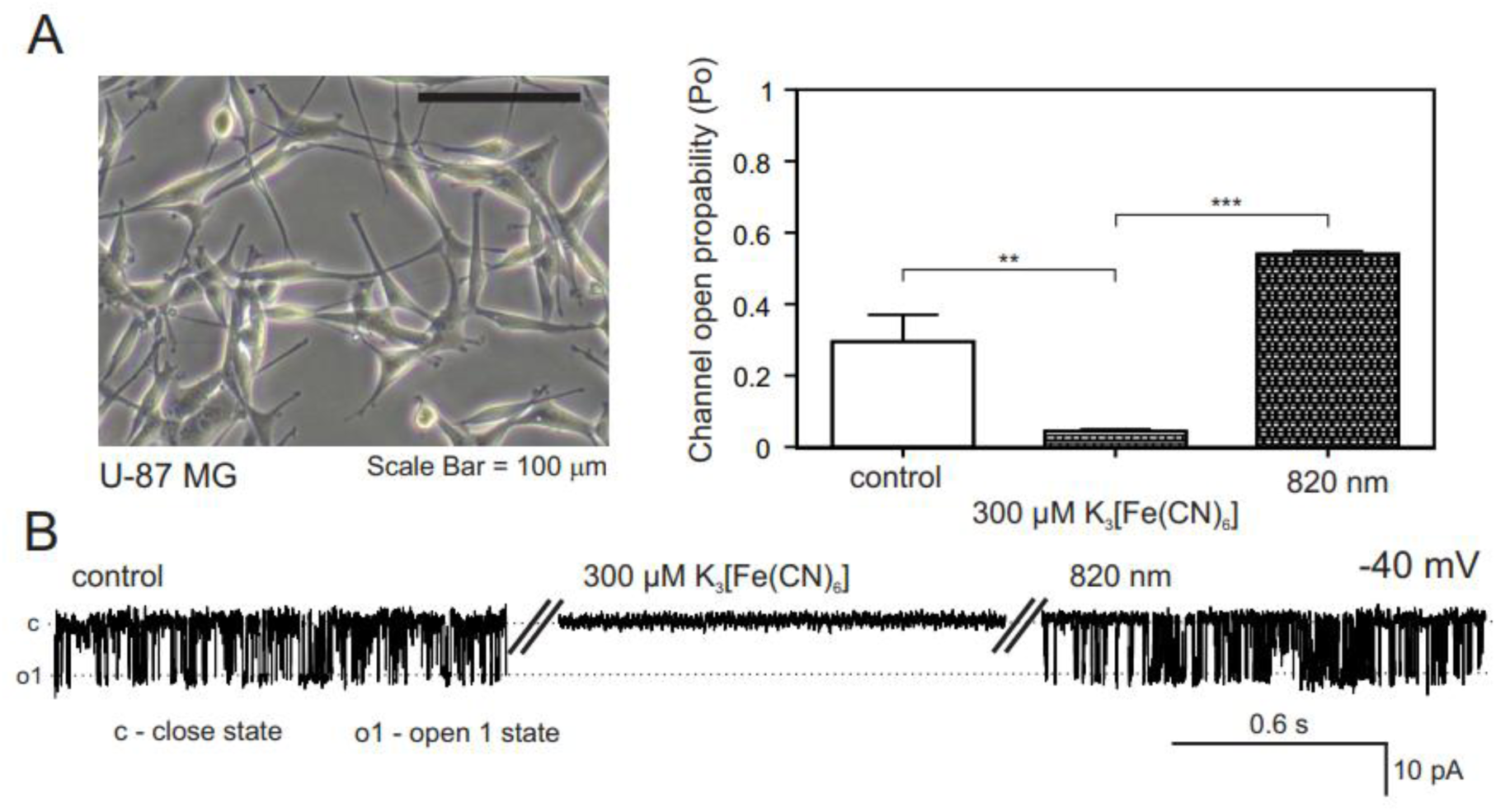
Regulation of the activity of BK_Ca_ channel in the mitoplast isolated from U-87 MG cells by NIR light. (A) U-87 MG cells illustrated with the light microscope. (B, C) Similarly to the hippocampal mitoplast, activity of BK_Ca_ channel in mitoplasts isolated from glioma cells significantly decreases after admission of 300 μM K_3_[Fe(CN)_6_] and is restored after 820 nm light illumination. Currents were recorded at −40 mV.

### 3.4. Changes in the transcriptome

To further characterize cellular responses to NIR illumination, we performed transcriptomic analysis of U-87 MG wild type (wt) and BK_Ca_-knockout (dBK) cells. Illumination resulted in differential expression of similar numbers of genes in both wt and dBK cells (2416 vs. 2358, respectively, Fig. 4), indicating global transcriptional response. Interestingly, the number of differentially expressed genes (DEGs) common to two groups was higher between two illuminated groups of cells (wt and dBK) than between treated and untreated dBK cells. The direction of changes was consistent between illuminated wt and dBK cells. However, 391 DEGs showed opposite regulation when comparing illuminated vs. non-treated wt cells and dBK_Ctrl vs. wt_Ctrl DEGs juxtapositions, suggesting that BK_Ca_ channel expression influences a subset of transcriptional responses. Reactome pathway analysis identified 51 and 72 significantly enriched pathways in illuminated wt and dBK cells, respectively. To minimize redundancy, pathways with fully overlapping DEG sets were excluded from further analysis. Six pathways were significantly enriched in illuminated wt cells but not in the illuminated dBK cells (Fig. 5, Tab. S1). Genes associated with the calnexin/calreticulin cycle pathway were down-regulated (Fig. 5A), whereas genes involved in ‘DDX58/IFIH1-mediated induction of interferon-α/β’ and ‘negative regulation of MAPK pathway’ were up-regulated. It seems worth underlining that DEGs from all those 6 pathways accounts for 19-35% of the total pool of genes assigned to those pathways (Fig. 5B, Tab. S1). Together, these results indicate that NIR illumination induces broad transcriptomic remodeling, while BK_Ca_ channels modulate specific components of this response.

**Fig. 4.**
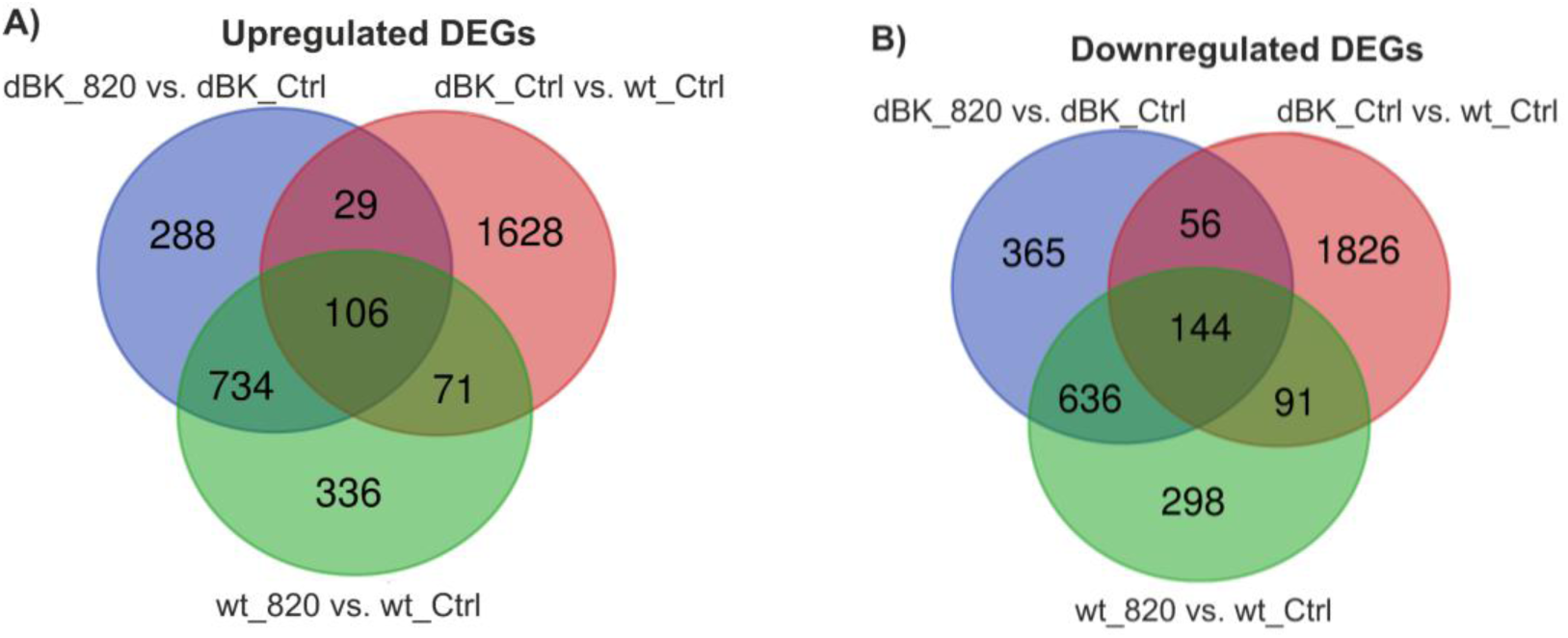
Venn diagrams of the differentially expressed genes (DEGs). A) and B) Diagrams drawn for up- and downregulated genes, respectively. ‘dBK_Ctrl vs. wt_Ctrl’ - indicates genes expressed differentially between unilluminated dBK and wt U-87 cells, respectively; ‘dBK820 vs. dBK_Ctrl’ - indicates genes expressed differentially between dBK U-87 cells illuminated with 820 light and unilluminated; ‘wt_820 vs. wt_Ctrl’ - indicates genes expressed differentially between wt U-87 cells illuminated with 820 light and unilluminated.

**Fig. 5.**
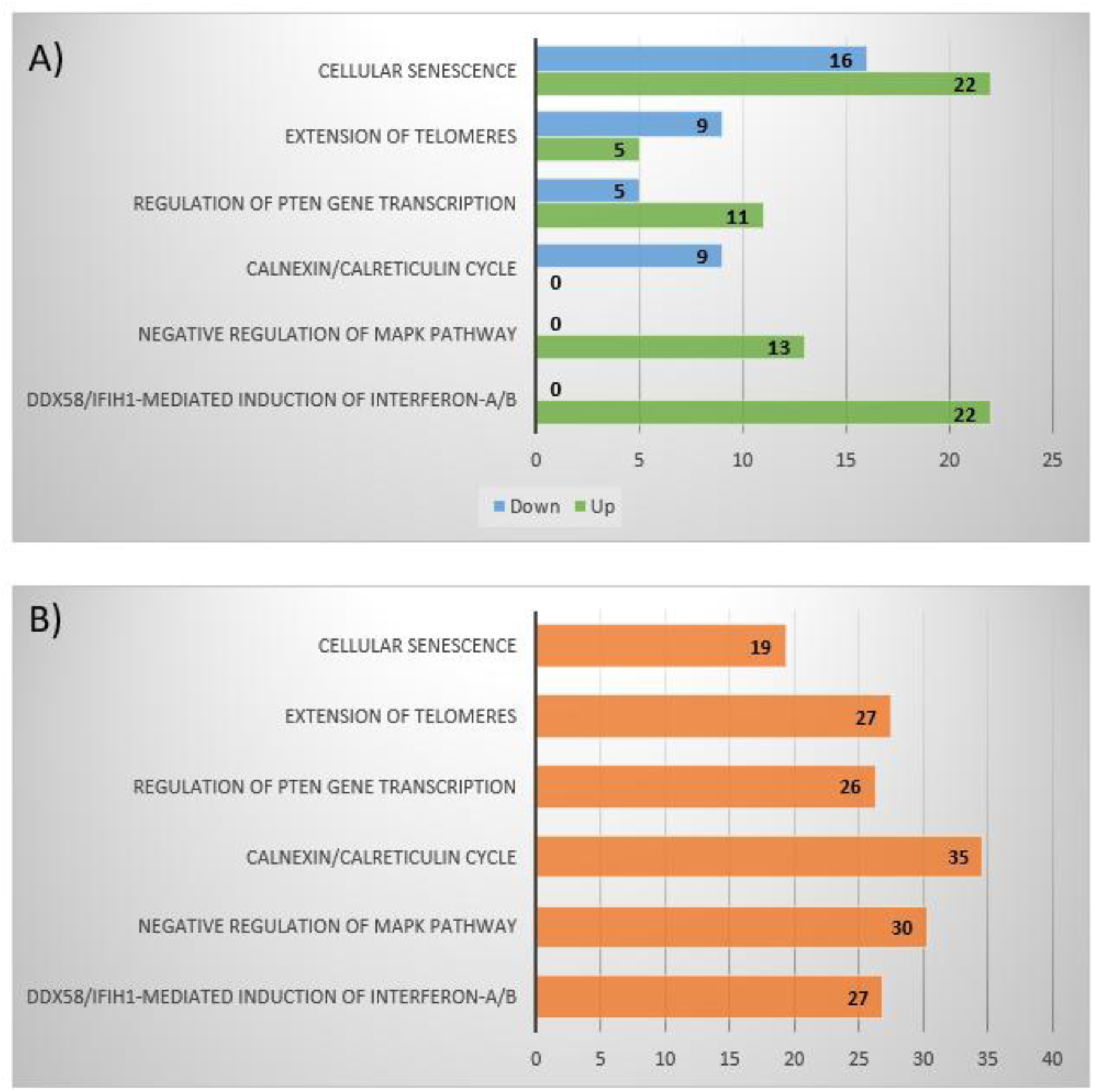
Reactome pathway analysis results. Pathways significantly enriched in U-87 dBK but not in the wt cells after illumination. The number of up and down-regulated differentially expressed genes from each pathway (A) and percentage of genes with changed expression levels compared to all genes assigned to the pathway (B) has been depicted. Pathways with DEGs contained in the list of DEGs from other pathways were omitted.

#### 3.4.1. DEGs encoding proteins localized in mitochondria (mitoDEGs)

To further focus on mitochondrial-related changes, we focused our analysis on differentially expressed genes encoding mitochondrial proteins (mitoDEGs). We identified 76 and 89 upregulated mitoDEGs in illuminated wt and dBK cells, respectively (Tab. S2), of which 45 were common to both types of cell. More mitoDEGs were downregulated after illumination - 149 and 123 in wt and dBK cells, respectively; with 77 mitoDEGs common to both groups. Fig. 6A and 6B illustrate mitoDEGs with 0.67>Fold-Change>1.33. Consistent with the overall DEG analysis, a significant part of mitoDEGs was common between illuminated wt and dBK cells, indicating a largely common mitochondrial response to NIR. Some mitoDEGs belong to the DEGs with the highest expression level changes (Fig. 6C, D). Functional analysis revealed that mitoDEGs encode proteins involved in mitochondrial dynamics and surveillance, mtDNA maintenance and mitochondrial translation, as well as metabolic processes (Fig. 7). Three genes show more than 2-fold expression increase in illuminated wt cells, whereas their expression was not significantly changed after illumination of dBK cells. Those are: STAR, BCO2 and MGARP, up-regulated in wt830vs.wtCtrl comparison with Fold-Change (FCh) of 26.84; 2.97 and 2.83, respectively. Moreover, overexpression of 4 genes with FCh>2 in this comparison (BCL2L11, NDUFC2, DNLZ and NAT8L) was larger than in dBK_830 vs. dBK_CTRL ratio by 47-75% and 4.79 fold in NAT8L case. Some of these genes encode key regulatory proteins in a metabolic pathway or core structural subunit of respiratory chain complex I. These results indicate that changes in the expression of mitochondrial genes are an indispensable component of the cellular response to PBM.

**Fig. 6.**
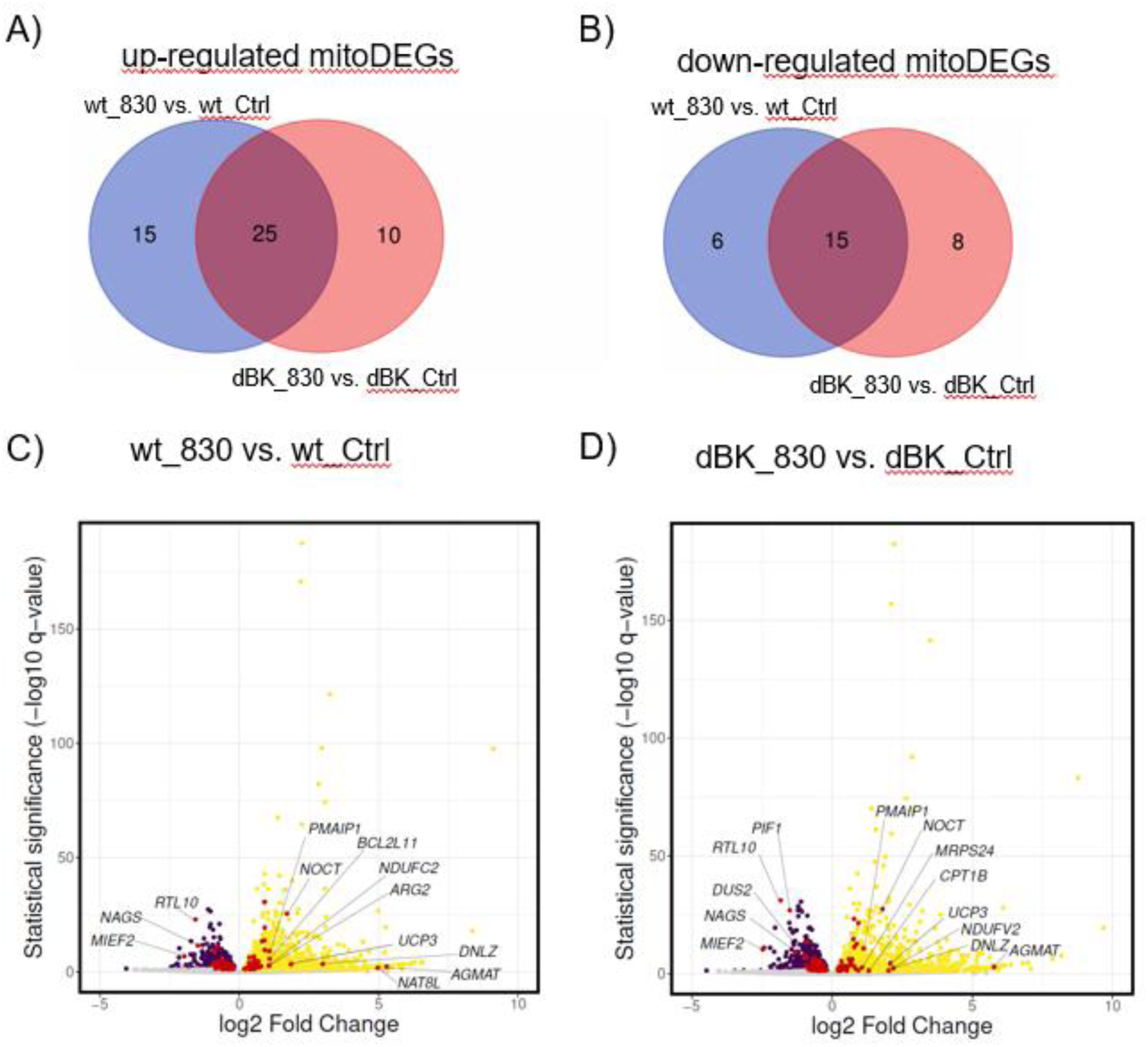
Genes with altered expression following exposure to near-infrared radiation of wt and dBK U-87 cells, encoding the human mitochondrial proteome (mitoDEGs). A) and B) Venn diagrams drawn for up- and downregulated mitoDEGs, respectively. Only mitoDEGs with 0.67>Fold-Change>1.33 were plotted. C) and D) volcano plots of mitoDEGs (red) from wt_820 vs. wt_Ctrl and dBK_820 vs. dBK_Ctrl comparison on the background of all up and down-regulated DEGs (yellow and violet, respectively). Only DEGs with q-value<0.05 were coloured. MitoDEGs with 1<log_2_ Fold-Change<-1 were labelled by Gene Symbol. ‘dBK820 vs. dBK_Ctrl’ and ‘wt_820 vs. wt_Ctrl’ - comparisons described in Fig. 4 legend.

**Fig. 7.**
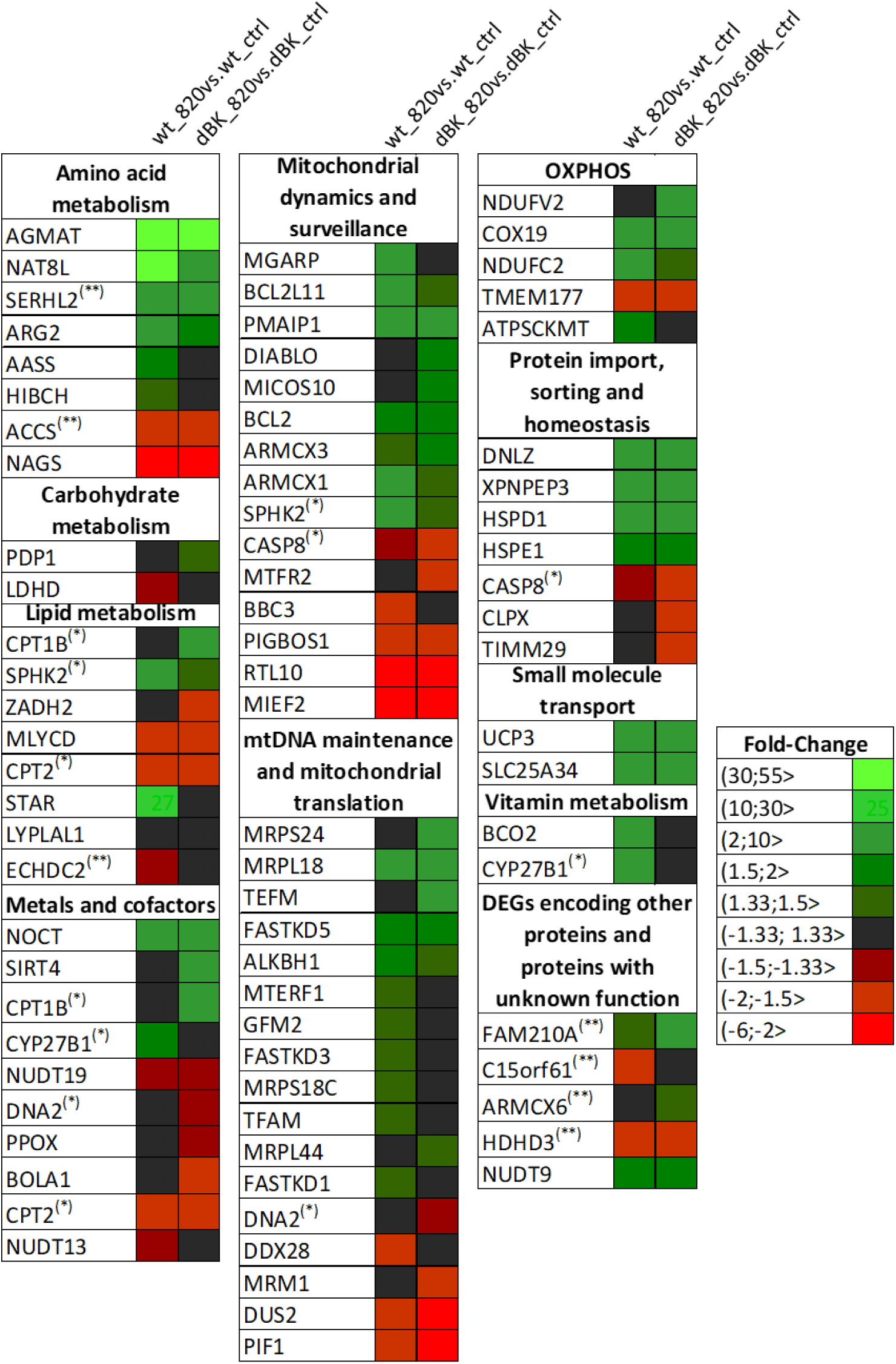
Functional characteristics of selected mitoDEGs. Differentially expressed genes encoding the human mitochondrial proteome (mitoDEGs) were divided into functional groups based on MitoPathways data available from MitoCarta database. Only mitoDEGs with 0.67>Fold-Change>1.33 in at least one comparison were plotted. (*) - mitoDEGs assigned to more than one functional group; (**) - function of the mitoDEGS is based on literature search due to lacking data in the mitoCarta. ‘dBK820 vs. dBK_Ctrl’ and ‘wt_820 vs. wt_Ctrl’ - comparisons described in Fig. 4 legend.

## 4. Discussion

Mitochondria host a diverse set of ion-conducting pathways, which are recognized as critical regulators of cellular bioenergetics including life and death decisions [34, 35]. The results presented here indicate that NIR light can regulate the activity of the mitoBK_Ca_ channel. The first evidence showing that BK_Ca_ channel is present at the inner mitochondrial membrane was given by Siemen and coworkers [36]. Those experiments were performed on the mitoplasts of the human glioma cell line [36, 37]. Since then, mitoBK_Ca_ channels were reported also in the mitoplast of rat astrocytes, organotypic hippocampal slice cultures and whole brain measured in lipid bilayers [22, 37–41] and functionally characterized [27]. Moreover, other potassium channels have also been identified in mitochondria derived from brain tissue. The Kv1.3 channel was found in mitochondria isolated from gerbil hippocampus and Kv 7.4 was found in mouse primary cortical neurons [42, 43]. The mitoK_ATP_ channel was first identified in brain mitochondria from glial and neuronal tissue of the rat cerebral cortex [44].

Both the mitoK_ATP_ and mitoBK_Ca_ channels mediate neuronal ischaemic preconditioning [45]. On the other hand, mitochondrial potassium channels are also potential targets in glioma treatment [46, 47]. Previously we reported functional coupling of the mitoBK_Ca_ channel to cytochrome c oxidase (COX), which is the main absorber of red and NIR light [2, 19]. This prompted questions if mitoBK_Ca_ channel is light-regulated and if neuroprotection by PBM can be optimized based on known mechanism of action. Ferricyanide (K_3_Fe(CN)_6_) can be used to oxidize COX and cytochrome c [48, 49]. It accepts electrons from components of the respiratory chain, mainly from cytochrome c and cytochrome bc1, oxidize cytochrome c [49] and COX [50]. To investigate role of COX in the effects of light on mitochondrial potassium channel, we decided to use K_3_Fe(CN)_6_ to model suppression of COX activity and 820 nm illumination to see if it can restore channel activity in our experimental model.

Our results suggest that 820 nm illumination can stimulate mitoBK_Ca_ channel opening, which makes this channel a possible responsive element of the intracellular PBM response. Although mitoBK_Ca_ channel does not contain chromophore itself, its regulation by the light is very quick, which suggests that it may participate in the acute response to PBM. Recently we published first to our knowledge data describing the influence of the light on mitoBK_Ca_ channel activity [20].

PBM treatment can offer significant improvement of many neurological diseases with no reported side effects [51–57]. Previous study reported, that that pulsed light therapy reversed the morphological changes and protein modifications caused by oxygen and glucose deprivation in OHC by attenuating inflammatory mechanisms [58]. Our study showed that mitoBK_Ca_ channel NS1619 and 820 nm light provides similar neuronal protection (Fig. 2). It is thus possible that this attenuation may have resulted, at least in part, from the activation of the mitoBK_Ca_ channel and reduced synthesis of reactive oxygen species (ROS). Our study suggests that PBM may be effective in preventing neuronal damage, and experiments conducted by Gerace and colleagues [58] also demonstrate the potential of this method as a post-injury treatment.

Another study performed on hippocampal cell line (HT-22) and mouse OHC assessed the influence of PBM on the oxidative stress in the hippocampus [59]. In both models illumination with the red LED light exerted an antioxidant effect. Moreover, this illumination inhibited cell death in HT-22 cells after H_2_O_2_ insult. Additionally, other studies on whole-animal and embryonic hippocampal culture illumination showed positive effects of PBM on mice models of epilepsy [60, 61].

Interestingly, even illumination of mice dorsum and hindlimbs with red light also showed neuroprotective potential for midbrain tyrosine hydroxylase-positive dopaminergic cells and FOS-positive neurons [62]. Furthermore, transcranial PBM with 1064 nm light-emitting diodes was also described as providing remarkable therapeutic effects in traumatic brain injury in mice [63].

Recent studies have demonstrated that near-infrared light modulates mitochondrial function by partially inhibiting cytochrome c oxidase activity, thereby limiting mitochondrial hyperpolarization, ROS generation, mitochondrial swelling, and excessive mitophagy during reperfusion injury [64]. These effects resemble several well-established consequences of mitoBK_Ca_ channel activation, including mild mitochondrial depolarization and attenuation of oxidative stress.

Influence of quick PBM on the transcriptome of human neuroblastoma SH-SY5Y cells was previously described at seven post-treatment time points [65]. Authors described that early phase (up to 8 h post-PBM) and late phase (24 h post-PBM) response shows different patterns of gene expression. That shows that changes following even short PBM last for many hours providing long-term response. Their findings also provide support for the hypothesis that PBM activates endogenous stress response pathways that in turn enhance cellular resilience. Based on the combined analysis of all seven post-treatment time points, 2647 genes were identified as differentially expressed. In accordance with our results, PBM of SH-SY5Y cells caused enrichment of various pathways related to the functions of the mitochondria and sumoylation pathway [65].

Study by Masha and coworkers screen for differential transcription only 84 genes involved in electron transport chain and oxidative phosphorylation [66]. uman fibroblasts modeling injury, diabetes, or hypoxia were shortly illuminated with a red diode laser. It resulted with upregulation of group-specific OXPHOS-related genes - coding for subunits of enzymes involved in complexes I, IV and V.

Recent study by Stevens et al. described transcriptomic changes after photobiomodulation in spinal cord injury [67]. Illumination with a red light resulted in downregulation of genes involved in electron transfer activity, COX activity and apoptotic process. Those results are consistent with a theory that slowing down oxidative phosphorylation can decrease the amount of ROS synthesized in the mitochondria and probability of cell death [23, 68–71]. The same mechanism of action is proposed for mitoK channels activators in ischemia-reperfusion (I/R) injury. Our study reports significant changes in expression of 35 OXPHOS-related genes (Tab. S2). Similarly to both described above experiments, the greatest changes were observed in subunits of complex I and assembly factors of complex IV and V. Furthermore, our data indicate significantly altered expression of 11 mitoDEG genes involved in apoptosis. Transcription levels of three of these genes decreased after illumination in cells expressing BK_Ca_ channels, but not in cells where the expression of these channels was inhibited. This suggests that their involvement as a secondary messenger in light-induced signaling may be BK_Ca_ channel-dependent. Down-regulation of all those 3 genes (ENDOG, BAD and HTRA2) in wt cells should cause an anti-apoptotic effect [72–74].

Most of the pathways revealed in our study as significantly changed in illuminated cells, but only in the presence of the BK_Ca_ channel, correlated with previously known functions of these channels. BK_Ca_ channel activity was shown to cause cytoprotection during I/R injury in cardiac and neuron tissue [75–77]. Its mechanism of action is associated with changes in inner mitochondrial membrane potential, mitochondrial Ca^2+^ influx, respiratory chain activity, and reactive oxygen species (ROS) synthesis [76, 77]. Pathways designated as ‘Negative regulation of MAPK pathway’ and ‘Regulation of PTEN gene transcription’ are also connected to apoptosis regulation [78–81]. The abovementioned mechanism shows the role of BK_Ca_ channel in cellular stress response [82–84]. Similarly, pathways associated with MAPK, PTEN; chaperons from ‘Calnexin/calreticulin cycle’; ‘cellular senescence’ or immunology-related ‘DDX58/IFIH1-mediated induction of interferon-α/β’ pathways take part in cellular signaling or response to stress-inducing factors [78, 85–89]. Moreover, PTEN and BK_Ca_ channels are both involved in cell growth, volume and size regulation [90, 91]. BK_Ca_ channel; ‘Negative regulation of MAPK pathway’ and ‘Extension of Telomeres’ pathway - all take part in control of the cell proliferation [81, 92–94].

In our study 22 and 16 genes from the ‘Cellular Senescence’ pathway were up- and downregulated, respectively, when wt cells were illuminated. It constitutes almost 20% of all genes assigned to this pathway by the Reactome Pathway Analysis tool. Interestingly in recent years it was reported that the level of BK_Ca_ pore-forming α subunit protein decreases in senescent cells with no detectable activity of this channel in the mitoplasts [95, 96].

In our current study, changes in activity of mitoBK_Ca_ channels can be observed immediately after NIR light illumination (Fig. 1, Fig. 3C). The activity of potassium channels directly affects the electrochemical gradient across mitochondrial inner membrane and their opening leads to rapid depolarization of mitochondrial membrane potential [19]. Meanwhile, most genes in mammalian cells are transcribed in bursts with timescales of tens of minutes to hours [97]. Moreover, transcriptomic profile after photobiomodulation changes at least between 15 min and 24 h after illumination [65]. Those data show that mitoBK_Ca_ channel is part of the acute response to NIR light, whereas transcriptomic and proteomic changes, along with cytoprotection, are observed as long-term effects.

In summary, we showed that the activity of the mitoBK_Ca_ channel can be restored by NIR light, both in mitoplast isolated from hippocampi and from U-87 MG cells. NIR light demonstrated cytoprotective effects in an ex vivo neuronal damage model. In the experimental setup used, the mitoBK_Ca_ channel activator provided an even higher level of cell protection, however its delivery and safe therapeutic dosing may be more challenging. Analysis of the transcriptome of NIR illuminated U-87 MG cells expressing and lacking BK_Ca_ channels showed thousands of genes with changed expression level. These genes enriched pathways associated with life and death decisions and some of them were dependent on BK_Ca_ channels expression (Tab. S1). The expression of the gene itself caused differential levels of mRNAs encoding hundreds of genes after illumination. Those genes belong to the metabolic pathways related to the BK_Ca_ channels functions. All those results indicate that BK_Ca_ channels play a role in acute and long-term effects of photobiomodulation and their activity can be regulated by NIR light.

## CRediT authorship contribution statement

Adam Szewczyk: Funding acquisition, Conceptualization, Supervision, Writing – review and editing. Barbara Zabłocka: Conceptualization, Supervision, Writing – review and editing. Piotr Bednarczyk: Conceptualization, Investigation, Data curation, Visualization, Writing – original draft, review and editing. Małgorzata Beręsewicz-Haller: Methodology, Investigation, Data curation, Visualization, Writing – original draft, review and editing. Joanna Lewandowska: Investigation, Data curation, Writing – review and editing. Bogusz Kulawiak: Conceptualization, Investigation, Data curation, Writing – review and editing. Antoni Wrzosek: Conceptualization, Investigation, Writing – review and editing. Barbara Kalenik: Methodology, Investigation, Data curation, Visualization, Writing – original draft, review and editing

## Declaration of generative AI and AI-assisted technologies in the manuscript preparation process

During the preparation of this work Joanna Lewandowska and Barbara Kalenik used ChatGPT-o4 in order to filter differentially expressed genes (DEGs) described in mitoCarta. Moreover, Barbara Kalenik used ChatGPT-4o in order to check if the list of DEGs from one pathway is included in the other pathway and ChatGPT-5.3-mini to language editing. After using this tool/service, the authors reviewed and edited the content as needed and took full responsibility for the content of the published article.

## Data availability

The datasets supporting the conclusions of this article are available in the GEO repository, accession number GSE325028.

